# When the tap runs dry: The multi-tissue gene expression and physiological responses of water deprived *Peromyscus eremicus*

**DOI:** 10.1101/2024.01.22.576658

**Authors:** Danielle M. Blumstein, Matthew D. MacManes

## Abstract

The harsh and dry conditions of desert environments have resulted in genomic adaptations, allowing for desert organisms to withstand prolonged drought, extreme temperatures, and limited food resources. Here, we present a comprehensive exploration of gene expression across five tissues (kidney, liver, lung, gastrointestinal tract, and hypothalamus) and 19 phenotypic measurements to explore the whole-organism physiological and genomic response to water deprivation in the desert-adapted cactus mouse (*Peromyscus eremicus*). The findings encompass the identification of differentially expressed genes and correlative analysis between phenotypes and gene expression patterns across multiple tissues. Specifically, we found robust activation of the vasopressin renin-angiotensin-aldosterone system (RAAS) pathways, whose primary function is to manage water and solute balance. Animals reduce food intake during water deprivation, and upregulation of *PCK1* highlights the adaptive response to reduced oral intake via its actions aimed at maintained serum glucose levels. Even with such responses to maintain water balance, hemoconcentration still occurred, prompting a protective downregulation of genes responsible for the production of clotting factors while simultaneously enhancing angiogenesis which is thought to maintains tissue perfusion. In this study, we elucidate the complex mechanisms involved in water balance in the desert-adapted cactus mouse, *P. eremicus*. By prioritizing a comprehensive analysis of whole-organism physiology and multi-tissue gene expression in a simulated desert environment, we describe the complex and successful response of regulatory processes.

## Introduction

Genomic adaptations play a pivotal role in enabling life to persist in the harsh and dynamic conditions of desert environments. Evolutionary processes have shaped the genomes of these organisms to enhance their capacity to withstand prolonged drought, extreme temperatures, and limited food resources (Colella et al., 2021; Tigano et al., 2022, 2020; Wu et al., 2014; Yang et al., 2016). Understanding these genetic underpinnings of desert adaptation not only contributes to our comprehension of evolutionary biology, but it also holds promise for insights into how these adaptations may be leveraged to address challenges posed by water scarcity and climate change in other organisms, including humans, and in other ecosystems. Studies of desert mammals have provided evidence of positive selection on genes related to food storage (Jirimutu et al., 2012; Wu et al., 2014), water reabsorption (Jirimutu et al., 2012; Marra et al., 2014, 2012; Yang et al., 2016), osmoregulation (Colella et al., 2021; Kordonowy and MacManes, 2017, 2016; MacManes and Eisen, 2014), fat metabolism (Chebii et al., 2020; Colella et al., 2021; Kim et al., 2016; Sugden et al., 2018; Tigano et al., 2020), thyroid-induced metabolism (Malaspinas et al., 2016), and salt regulation (Ababaikeri et al., 2020). These genetic insights suggest the molecular basis of observed phenotypes, including enhanced metabolic water production (Frank, 1988; MacMillen and Hinds, 1983; Walsberg, 2000), reduced water loss (Blumstein and MacManes, 2023; Frank, 1988; Schmidt-Nielsen, 1975), tolerance to high-salt diets (Ali et al., 2019; Jirimutu et al., 2012), and coping with starvation and dehydration (Blumstein and MacManes, 2023, 2023; Boumansour et al., 2021; Kordonowy et al., 2017; Kordonowy and MacManes, 2017; MacManes, 2017), which are all common in desert-dwelling mammals. However, it remains unclear how water deprivation affects the activities of other organs with respect to their expression of genes in the whole-organism context.

Osmoregulation, or the process through which animals manage water and solute balance, is critical for desert animals. It involves the maintenance of internal fluid homeostasis, with water intake being dependent on factors such as drinking, dietary sources, and metabolic water, while water output is regulated through processes including waste removal (i.e., urine and feces), respiration, perspiration, and reduced food intake (Bouby and Fernandes, 2003; Popkin et al., 2010; Watts and Boyle, 2010). Failure to maintain water and solute homeostasis can result in impaired renal, reproductive, and cardiovascular function, affect an animal’s ability to regulate its body temperature, and ultimately lead to death (Popkin et al., 2010). During water deprivation, a decrease in extracellular water volume results in heightened plasma osmolality due to an elevated concentration of solutes, primarily sodium, is detected by osmoreceptors (Greenleaf, 1992; Leib et al., 2016; Thornton, 2010). Osmotic balance is then intricately managed through thought two distinct multisystem mechanisms, the renin-angiotensin-aldosterone system (RAAS) and by vasopressin (Aisenbrey et al., 1981; Bouby and Fernandes, 2003; Greenleaf, 1992; Roberts et al., 2011).

In response to changes in osmotic pressure, osmoreceptors in the hypothalamus become activated, aldosterone is released from the adrenal glands, and vasopressin is produced in the hypothalamus and released from the posterior pituitary gland (Leib et al., 2016; Thornton, 2010; Yoshimura et al., 2021). This promotes water reabsorption in the kidneys by enhancing the water permeability of epithelial cells lining renal collecting ducts (Fuller et al., 2020). Additionally, the kidneys retain sodium while also excreting solutes via urination (Qian, 2018). Proteins related to the transport of water are translocated from within the cell to the cell surface, forming water channels, resulting in increased water reabsorption from the tubule system of the nephrons back into the bloodstream (Brown et al., 1995; Kortenoeven and Fenton, 2014; Verkman, 2002) and the retention of sodium in the distal tubules (Goodfriend, 2006). Renin is released from the kidney, and acts on angiotensinogen that is released from the liver (Blair et al., 1997; Greenleaf, 1992). This triggers the formation of angiotensin I, which is then converted into angiotensin II, a hormone involved in regulating blood pressure and peripheral circulation, by the release of angiotensin-converting enzyme in the lungs (Fountain et al., 2023; Greenleaf, 1992; Santos et al., 2019). As a result, RAAS impacts water and solute retention by promoting vasoconstriction, stimulating the release of aldosterone, and enhancing the reabsorption of sodium and water in the kidneys. This orchestrated response helps regulate blood pressure and maintain fluid balance in the body.

If water homeostasis is not achieved, blood volume continues to decrease, resulting in hemoconcentration. As dehydration leads to a higher concentration of erythrocytes, its viscosity increases, potentially enhancing the risk of spontaneous clot formation and hindering blood flow through vessels. This elevated blood viscosity might impede the efficient delivery of oxygen and nutrients to various tissues and organs, triggering alterations in vascular dynamics in nutrient-deprived, hypoxic environments (Alonso et al., 2005; Cavaglia et al., 2001; Swain et al., 2003). Studies suggest that impaired flow-dynamics typical of dehydration may induce reversible changes in angiogenesis, altering the growth and development of blood vessels to regulate blood flow and distribution (Alim et al., 2019; Dumas et al., 2020). The modulation of angiogenesis during dehydration reflects the body’s dynamic response in adjusting vascular networks to manage the metabolic demands of tissues.

Vasopressin receptors expressed in the kidneys, lungs, liver, hypothalamus etc., further affect vasoconstriction, glycogenolysis, water reabsorption, thermoregulation, and food intake (Yoshimura et al., 2021). Water deprived animals have been shown to limit food intake (Armstrong et al., 1980, Blumstein, MacManes, personal observation) as an adaptive mechanism allowing for osmotically sequestered water in the GI to be reabsorbed into the systemic vasculature (Kutscher, 1968; Lepkovsky et al., 1957; Schoorlemmer and Evered, 1993) which thereby reduces solute load (Rowland, 2007). Decreased food intake may be driven by the expression of vasopressin in the hypothalamus (Yoshimura et al., 2021), suggesting another link between eating and drinking. To survive reduced food intake during water deprivation (i.e., dehydration anorexia) while maintaining blood glucose concentrations, rodents have been previous studies have shown increased glycogenolysis, lipolysis, and/ or gluconeogenesis (Salter and Watts, 2003; Schoorlemmer and Evered, 2002; Watts and Boyle, 2010).

In this this study, we have performed a comprehensive analysis of gene expression across five tissues relevant to the response to dehydration (kidney, liver, lung, gastrointestinal tract, and hypothalamus) and used 19 phenotypic measurements to assess the whole-organism physiological and genomic response to water deprivation in a hot and dry environment in the desert-adapted cactus mouse (*Peromyscus eremicus*). The results of this study, including: 1) a robust activation of RAAS, as seen by upregulation of *AGT* across all five tissues, 2) upregulation of *PCK1*, reflecting an adaptive response to maintain blood glucose levels during decreased oral intake, 3) a broad decrease in genes related to coagulation, possibly in response to hemoconcentration and 4) a clear signal of vascular remodeling. Overall, the lung experienced the largest number of changes in gene expression, followed by tissues involved in RAAS and then the hypothalamus. Each tissue differed in its own way with regards to the number of genes with highly correlated expression profiles; however, this was not due to different gene expression between the various tissues.

## Methods

### Animal Care, RNA Extraction, and Sequencing

Captive born, sexually mature, non-reproductive healthy male and female *P. eremicus* were reared in an environmental chamber designed to simulate the Sonoran desert (Blumstein et al., 2022; Blumstein and MacManes, 2023; Colella et al., 2021; Kordonowy et al., 2017). All mice were subjected to standard animal care procedures before the experiment which included a health assessment conducted by licensed veterinary staff following animal care procedures guidelines established by the American Society of Mammologists (Sikes et al., 2016) and approved by the University of New Hampshire Institutional Animal Care and Use Committee under protocol number 210602. Mice were provided a standard diet and fed *ad libitum* (LabDiet® 5015*, 26.101% fat, 19.752% protein, 54.148% carbohydrates, energy 15.02 kJ/g, food quotient [FQ] 0.89). Animals were randomly selected and assigned to the two water treatment groups (n=9 of each group, female mice with water, female mice without water, male mice with water, and male mice without water, total n=36). Prior to the start of the experiment, a temperature-sensing passive integrated transponder (PIT) tag (BioThermo13, accuracy ±0.5°C, BioMark®, Boise, ID, USA) was implanted subdermally. At the start of the experiment (day 0, time 0hr, 10:00), mice were weighed (rounded to the nearest tenth of a gram) on a digital scale and water was removed from chambers corresponding to those animals in the dehydration group. Mice were metabolically phenotyped for the duration of the experiment (Blumstein and MacManes, 2023) using a pull flow-through respirometry system from Sable Systems International (SSI). Rates of CO_2_ production, O_2_ consumption, and water loss were calculated using equations 10.6, 10.5, and 10.9, respectively, from Lighton (2018). Respiratory quotient (RQ, the ratio of VCO_2_ to VO_2_) and energy expenditure (EE) kJ hr^-1^ were calculated as in Lighton (2018, eq. 9.15). For downstream analysis, we calculated the mean of the last hour of water loss, EE, and RQ for each mouse.

At the conclusion of the experiment (day 3, time 72hr, 12:00) as described in Blumstein and MacManes (2023), body temperature was recorded via a Biomark® HPR Plus reader, mice were weighed, animals were euthanized with an overdose of isoflurane, and 120 µl of trunk blood was collected for serum electrolyte measurement and analyzed with an Abaxis i-STAT® Alinity machine using i-STAT CHEM8+ cartridges (Abbott Park, IL, USA, Abbott Point of Care Inc). We measured the concentration of sodium (Na, mmol/L), potassium (K, mmol/L), blood urea nitrogen (BUN, mmol/L), hematocrit (Hct, % PCV), ionized calcium (iCa, mmol/L), glucose (Glu, mmol/L), osmolality (mmol/L), hemoglobin (Hb, g/dl), chlorine (Cl, mEq/L), total CO_2_ (TCO_2_, mmol/L), and Anion gap (AnGap, mEq/L). Using Na, Glu, and BUN, we calculated serum osmolality. To test for statistically significant (p < 0.05) differences, we used a student’s two-tailed t-test (stats::t.test) between the sexes for each experimental group in R v 4.0.3 (R Core Team, 2020).

The lung, liver, kidney, a section of the large intestines (referred to as GI throughout), and hypothalamus were collected and stored in RNAlater (Ambion) at 4°C for 12hr before being frozen at −80°C for long-term storage. Prior to RNA extraction, the tissues were removed from the RNAlater and a small section was dissected off. Care was taken to retain an anatomically similar region of tissue from each animal. Tissues were mechanically lysed using a Bead Beater, and RNA was then extracted using a standardized Trizol protocol. RNA libraries were prepared using standard poly-A tail purification, prepared using Illumina primers, and individually dual-barcoded using a New England Biolabs Ultra II Directional kit (NEB #E7765). Individually barcoded samples were pooled and sequenced paired end and 150 bp in length on two lanes of a Novaseq at the University of New Hampshire Hubbard Center of Genome Studies.

### Genome Alignment and Differential Gene Expression

All the code used to analyze the data are located at the GitHub repository (https://github.com/DaniBlumstein/dehy_rnaseq). The *P. eremicus* genome version 2.0.1 from the DNA Zoo Consortium (dnazoo.org) was indexed, and reads from each individual were aligned to the genome using STAR version 2.7.10b (Dobin et al., 2013), allowing a 10 base mismatches, a maximum of 20 multiple alignments per read, and discarding reads that mapped at <30% of the read length. Aligned reads were counted using HTSEQ-COUNT version 2.0.2 (Anders et al., 2015).

Counts from HTSEQ-COUNT were exported as csv files, and all downstream statistical analyses were conducted in R v4.0.3 (R Core Team, 2020). Counts were merged into a gene-level count by combining all counts that mapped to the same gene. Low expression genes (defined as having 10 or less counts in 8 or more individuals) were removed from downstream analyses. Differential gene expression analysis was conducted in R using DESEQ2 (Love et al., 2014). For the dataset as a whole, we performed three models to test for the effects of sex, water access, and tissue type. For each tissue, we performed two models, testing the effect of and identifying genes specific to sex and water access with a Wald test. Results were visualized using GGPLOT2 (Wickham, 2016).

### Weighted Gene Correlation Network Analysis

To identify the regulation of gene expression associated with responses to water access, we performed a weighted gene correlation network analysis (WGCNA), a network-based statistical approach that identifies clusters of genes with highly correlated expression profiles (modules), (Langfelder and Horvath, 2008) for each tissue independently. This approach allows us to relate gene expression with physiological phenotypes (mean EE, water loss, RQ, total weight loss, proportional weight loss, sex, body temperature, water access, and the panel of electrolytes). Prior to WGCNA, read counts were normalized within tissues using DESEQ2 (Love et al., 2014). Module detection was done using WGCNA::blockwiseModules with networkType set to “signed” but otherwise default parameters were used. We estimated a soft threshold power (β) for each tissue dataset by plotting this value against mean connectivity to determine the minimum value at which mean connectivity asymptotes, which represents scale-free topology (liver = 15, kidney = 21, GI = 14, lung = 20, hypothalamus = 14).

### Canonical Correlation Analysis

We used a Canonical Correlation Analysis (CCA) implemented in the R package vegan (Oksanen, 2010) to investigate multivariate correlation of gene expression, by tissue, water access, and sex, with metabolic variables (mean EE, mean RQ, mean water loss, body temperature, and proportional weight loss) and display the three levels of information in a triplot. We used an ANOVA to identify what response variables were significant. Significant response variables were graphed as vectors and allowed us to identify their correlative nature; vectors pointing in the same direction are positively correlated, while vectors pointing in opposite direction are negatively correlated. To identify genes of interest, we selected genes that graphed two standard deviations away from the mean for CCA1 and CCA2.

### Gene Ontology

To examine gene ontology of DE genes and WGCNA modules, we cross-referenced our gene IDs with *Homo sapiens* gene IDs via Ensembl before running Gene Ontology (GO) analyses. Each analysis above resulted in a list or lists of genes that were used as input for the GO analysis using the R package gprofiler (Kolberg et al., 2023). From there, we identified the topmost 20 significant GO terms based on g:SCS corrected p-values (Reimand et al., 2007) for each up and downregulated list for each tissue and for each significant module from the individual WGCNA analysis.

### Consensus gene list and KEGG pathway analysis

We generated a high confidence consensus list of genes from the results of the three orthogonal analyses; DE, WGCNA, and genes located two standard deviations away from the origin in the CCA. We then selected three KEGG pathways (Renin-Angiotensin system KEGG pathway [hsa04614], vasopressin-regulated water reabsorption pathway [hsa04962], and insulin resistance KEGG pathway [hsa04931], Kanehisa et al., 2023; Kanehisa and Goto, 2000) based on genes in our consensus gene set and cross references the genes in those pathways with significantly differently expressed genes in our five tissue datasets.

## Results

No health issues were detected by veterinary staff, no animals were removed prior to the end of the experiment, and all mice were active at the end of the experiment.

### Genomic Data

We obtained an average of 21.44 million reads (+-11.6 million SD) per sample (PRJNA1048512). On average, 78.33% of reads were uniquely mapped per sample (+-2.12% SD). Data on the number of reads and mapping rate per sample are located in Supplemental File 1, raw read files are archived at NCBI SRA BioProject: PRJNA1048512, and all gene expression count data and code used to analyze the data are located at the GitHub repository (https://github.com/DaniBlumstein/dehy_rnaseq).

### Electrolytes and Physiological Phenotypes

The same mice used to generate the electrolyte and physiology data more fully described in Blumstein and MacManes (2023) are also used in the study described herein for RNAseq analysis. When comparing males and females separately, the following electrolytes showed significant differences with and without access to water: Na (male and female Na p = 0.0016 and p = 0.0026 respectively), BUN (p = 0.001/0.003), Hct (p = 0.002/0.001), osmolality (p = 8.2^-05^/0.0001), Cl (p = 0.02/0.007), Hb (p = 0.017/0.009), and TCO2 (female p = 0.017) (Table 1). When comparing males to females within each water treatment (with or without access to water), no significant differences were found in the electrolyte levels (Table 1). Both males and females experienced significant weight loss (p = 0.001, 0.005) and proportional weight loss (p = 2.2e-16, 2.659e-09) at the end of the experiment (Blumstein and MacManes, 2023). Body temperature was significantly lower for female mice without access to water, but not for males (p = 0.0003).

**Table 1.** Mean measurements for serum electrolyte measurements (Na = Sodium (mmol/L), K = Potassium (mmol/L), Cr = Creatinine (μmol/L), BUN = Blood Urea Nitrogen (mmol/L), Hct = Hematocrit (% PCV), iCa = Ionized Calcium (mmol/L), and osmolality (mmol/L), change in weight (g), and body temperature (°C) for female (n=18), male (n=18), and all *Peromyscus eremicus* (n=36) with and without access to water. Data were collected and are further described in Blumstein and MacManes (2023).

| sex | all |  | female |  | male |  |
| --- | --- | --- | --- | --- | --- | --- |
| water access | no | yes | no | yes | no | yes |
| Hct | 43.31 | 33.18 | 44.38 | 32.50 | 42.25 | 33.78 |
| Hb | 14.73 | 11.28 | 15.09 | 11.05 | 14.36 | 11.48 |
| Na | 155.75 | 144.47 | 159.25 | 144.75 | 152.25 | 144.22 |
| K | 6.58 | 5.88 | 6.19 | 5.85 | 6.98 | 5.91 |
| Cl | 123.63 | 115.59 | 125.88 | 115.38 | 121.38 | 115.78 |
| TCO<sub>2</sub> | 24.31 | 20.53 | 25.63 | 20.50 | 23.00 | 20.56 |
| BUN | 56.75 | 33.53 | 54.25 | 33.63 | 59.25 | 33.44 |
| Crea | 0.24 | 0.20 | 0.28 | 0.20 | 0.21 | 0.20 |
| Glu | 116.88 | 121.53 | 117.38 | 122.38 | 116.38 | 120.78 |
| iCa | 1.25 | 1.30 | 1.28 | 1.31 | 1.22 | 1.28 |
| AnGap | 15.19 | 15.18 | 14.50 | 15.38 | 15.88 | 15.00 |
| osmolality | 300.68 | 290.57 | 306.13 | 274.05 | 295.22 | 307.10 |
| change in weight | -5.07 | -0.03 | -4.75 | 0.08 | -5.38 | -0.15 |
| body temperature | 35.46 | 36.13 | 34.94 | 36.16 | 35.98 | 36.10 |

To relate whole-organism physiology data to gene expression data, we calculated the means for each mouse from the last hour of data collected in Blumstein and MacManes (2023, data located at: https://github.com/DaniBlumstein/dehy_phys) for WLR, EE, and RQ for the same 18 adult females and 18 adult males (n=9 of each treatment, total n=36) used in this study. Within sex, WLR significantly different between water groups (male and female, p = 0.001 and 0.002), RQ was significantly different between water groups for males (p=0.0003), and EE was not significantly different for either males or females.

### Differential Gene Expression

We cross-referenced our gene IDs with *Homo sapiens* gene IDs. Patterns of gene expression data are largely driven by tissue type (PC1: 42% variance and PC2: 26% variance, Supplemental Figure 1).

We then conducted all downstream analyses (except for CCA, see below) on each tissue independently. Count and sample data were filtered to include only the tissue of interest and low expression genes were removed. This resulted in removing 4929 genes in the kidney and left 12083, 4390 genes removed in the GI leaving 12622 genes, 3623 genes removed in the hypothalamus leaving 13389 genes, 5887 genes were removed in the liver leaving 11125 genes, and 4070 genes were removed in the lung leaving 12942 genes. Within each tissue, we found many differentially expressed genes (p < 0.05) between water treatments (Figure 1, Table 2) and few differentially expressed genes between sex (Table 2).

**Figure 1.**
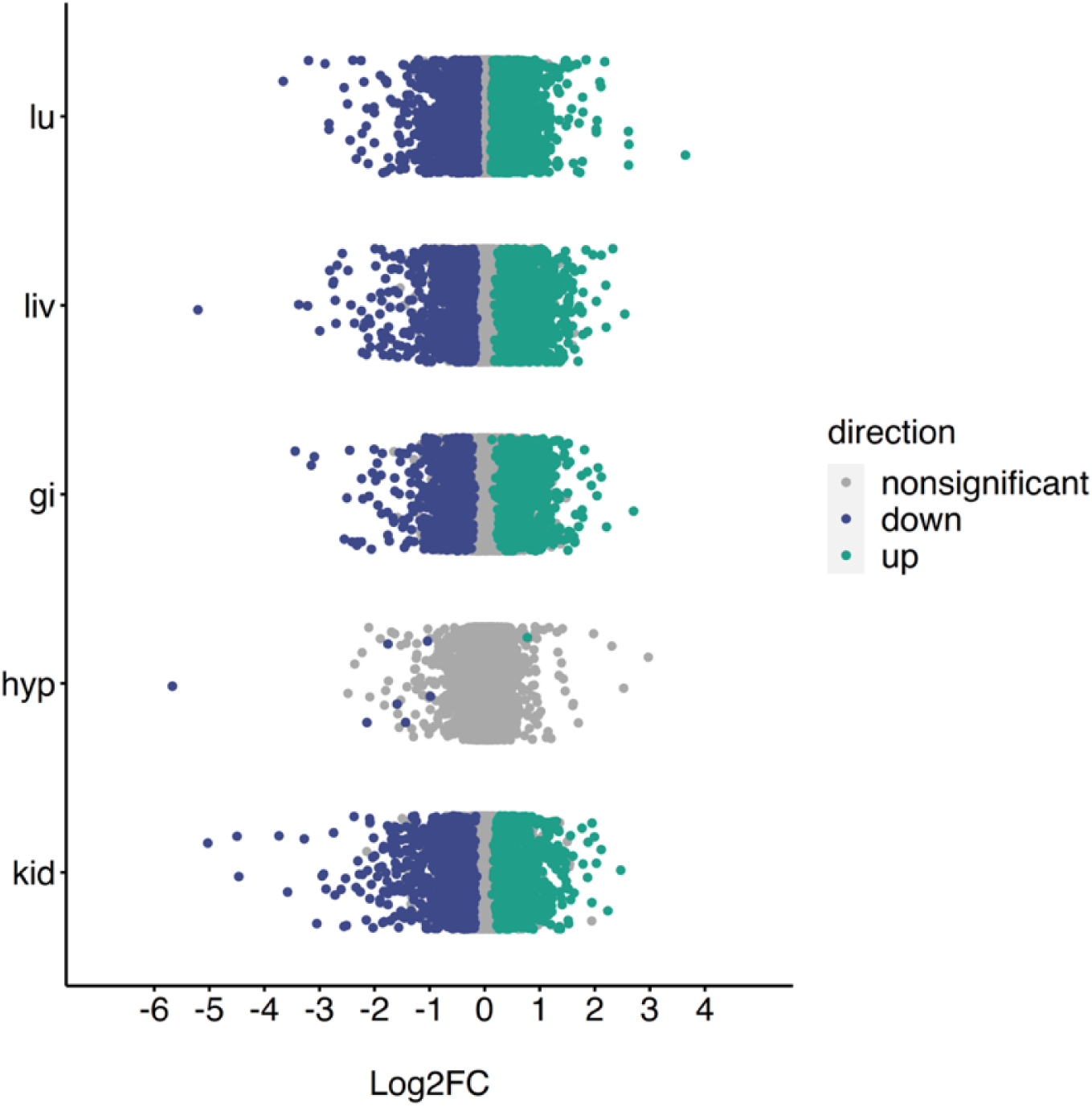
Log2FC of all genes across the lung (lu), liver (liv), gastrointestinal tract (gi), hypothalamus (hyp), and kidney (kid) of *Peromyscus eremicus* with water vs without water. Blue and green colored dots indicate p<0.05, whereas grey dots indicate p>=0.05.

**Table 2.** The number of differentially expressed (DE) genes in the lung, liver, gastrointestinal tract, hypothalamus, and kidney of *Peromyscus eremicus* for with water vs without water and males vs females (adjusted p-value < 0.05).

| Tissue | DE between water treatments<br>with water vs without water |  | DE between sexes<br>males vs females |  |
| --- | --- | --- | --- | --- |
|  | Up | Down | Up | Down |
| kidney | 793 | 1056 | 5 | 7 |
| liver | 933 | 1122 | 1 | 3 |
| lung | 2190 | 2183 | 5 | 5 |
| hypothalamus | 1 | 8 | 1 | 2 |
| gastrointestinal tract | 956 | 703 | 1 | 1 |

### Weighted Gene Correlation Network Analysis

A total of 12083 genes in the kidney were successfully assigned into 13 modules with the number of genes per module ranging from 31 – 7008. A full list of gene assignments is available in Supplemental Table 3. Of the 13 modules identified, 9 modules were significant for three or more phenotypes (Supplemental Table 2). We identified 24 individual modules using 12622 genes for the GI. The modules contained 33-3530 genes each (Supplemental Table 2). Of these modules, 14 were significantly correlated with three or more phenotypes (Supplemental Table 2). In the lung, 12942 genes were assigned to 13 modules. Each module contained 23-7116 genes (Supplemental Table 2). Nine modules were significant for three or more phenotypes (Supplemental Table 2). A total of 13389 genes were assigned to 18 different modules in the hypothalamus. Modules contained 30-2319 genes (Supplemental Table 2). Of the 18 modules, three modules were significant for three or more phenotypes (Supplemental Table 2). Finally, 11125 genes were assigned to 18 modules in the liver, with the number of genes per module ranging from 28-3116 (Supplemental Table 2). Of the 18 modules, 13 modules were significant for three or more phenotypes (Supplemental Table 2).

### Gene Ontology

After filtering to the top 20 GO terms for the upregulated and downregulated significantly differentially expressed genes for each tissue, we identified 149 unique terms. Of the 149 GO terms, 14 terms were identified in two different tissues (Supplemental Figure 2).

For each tissue WGCNA analysis, we identified GO terms for each significant module phenotype combination. After filtering each combination for the top 20 GO terms, we identified a total of 379 unique GO terms. 199 GO terms were identified in the hypothalamus with 168 terms only appearing in one module, 25 identified in two modules, five in three modules, and one in four modules (Figure 2). Within the kidney 82 unique GO terms were identified. Specifically, 78 terms were in one module and four terms were in two modules. There were 122 unique GO terms identified in the liver with 100 of the go terms appearing in one module, nine terms in two modules, and three terms in three different modules (Figure 2). Lastly, the lung had 52 unique GO terms identified, the fewest number any tissue, and there was no GO term overlap for any of the modules (Figure 2). When comparing GO terms between tissues, 53 terms were in two different tissues, five terms were in three of the tissues, and one term were in four tissues (Figure 2).

**Figure 2.**
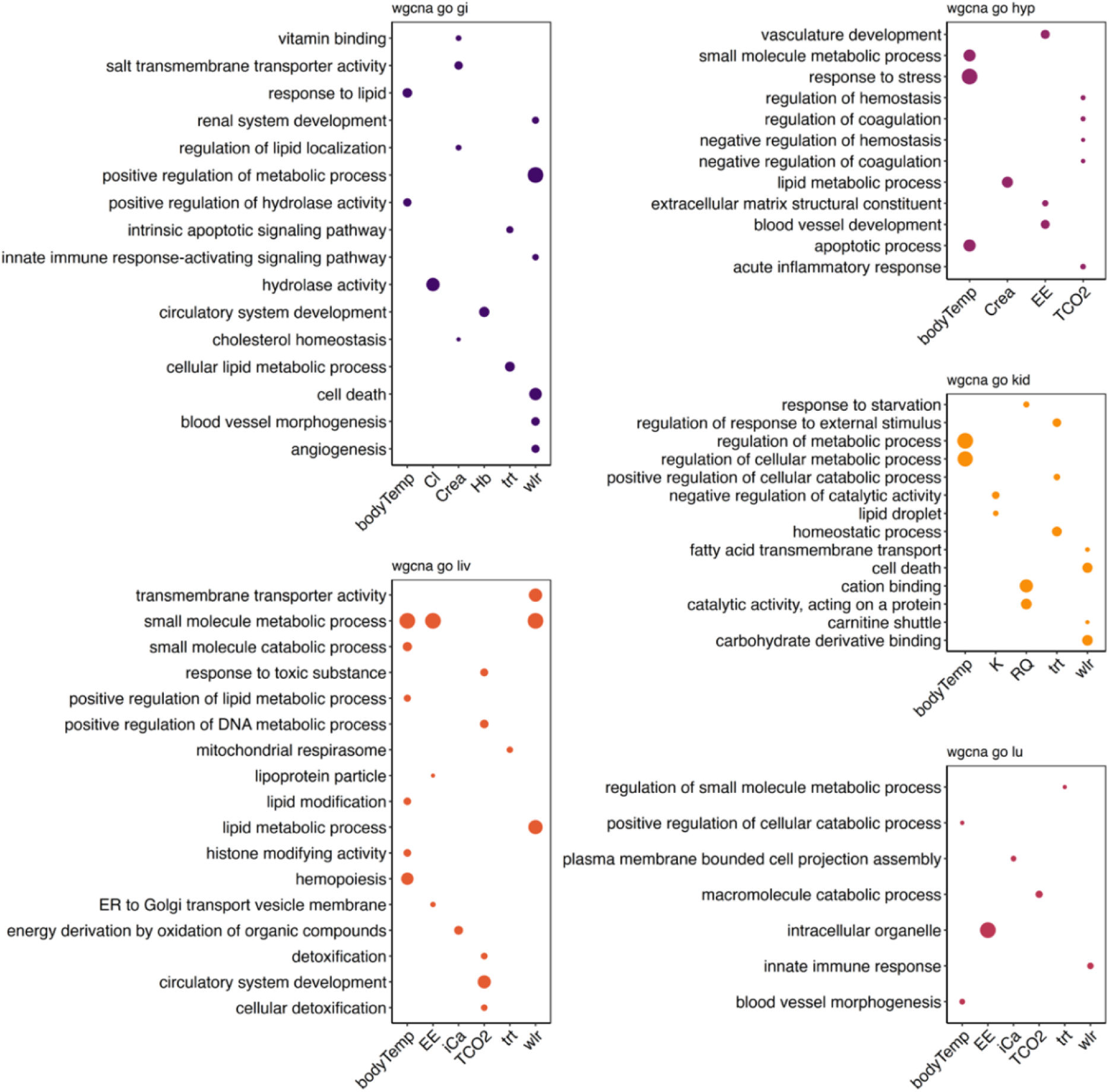
Visualization of gene ontology (GO) terms to show common WGCNA modules within and between the lung (lu), liver (liv), gastrointestinal tract (gi), hypothalamus (hyp), and kidney (kid) of *Peromyscus eremicus*. Visualized are selections of the top 20 significant GO terms for each phenotype module combination. The number of genes in the GO term are indicated by size of the dots.

### Canonical Correlation Analysis

We examined the relationship between gene expression and water access, physiological variables (EE, RQ, WLR, proportional weight loss, body temperature), and tissue type using CCA (Oksanen, 2010). CCA suggests a significant overall association between the physiological variables and gene expression across tissues (F = 105.45, p = 0.001; CCA1 – 38.02% and CCA2 – 21.14%, Figure 3, Table 4). We identified 1233 genes two standard deviations from the origin and found a high degree of overlap between the genes located two standard deviations in the CCA and genes assigned to WGCNA modules (kidney: 37/10906, GI: 60/11032, lung: 69/12527, hypothalamus: 53/1971, liver: 40/11051). Here, the proportional weight loss and WLR are significantly correlated (both p=0.001, Table 4) with gene expression in the hypothalamus and GI (Figure 3), but there is no overlap between the genes and the two vectors. This suggests that proportional weight loss and WLR exert different effects on gene expression in the hypothalamus and the GI.

**Figure 3.**
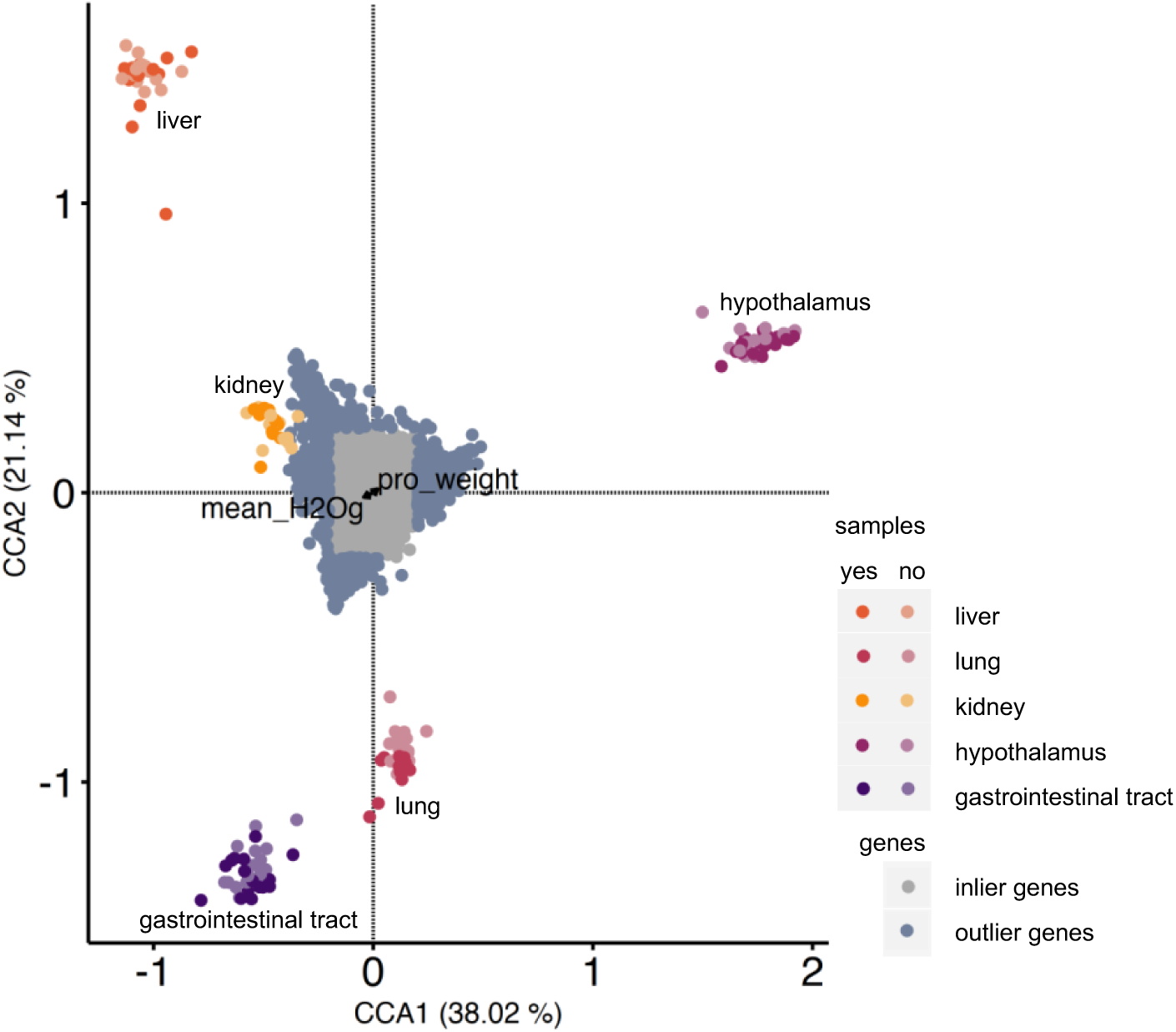
Canonical correspondence analysis (CCA) indicates correlations between normalized differentially expressed genes and physiological measurements for *Peromyscus eremicus* with and without access to water. The distribution of tissue samples in Euclidian space as a function of their gene expression values is shown (points colored by tissue type and water treatment). Outlier genes (defined as two standard deviations or more from the mean) are colored blue. Inlier genes (defined as less than two standard deviations from the mean) are colored grey. CCA reveals a significant relationship between proportion weight loss (F = 4.8006, P = 0.001) and, while not significant, a strong relationship for water loss rate (WLR) (*F* = 1.9856, *P* = 0.096). This can be seen by a subset of genes (blue) pulled in the direction of the proportion weight loss (pro_weight) ordination vector and a subset of genes (blue) pulled in the direction of the WLR (mean_H_2_Og) ordination vector.

**Table 4.** ANOVA results from the Canonical Correspondence Analysis. Formula: gene expression ∼ water access + proportional weight loss + RQ + EE + WLR + body temperature + sex + tissue).

|  | Df | ChiSquare | F | Pr(>F) |
| --- | --- | --- | --- | --- |
| water access | 1 | 0.00012 | 6.437 | 0.001 |
| proportional weight loss | 1 | 0.00009 | 4.801 | 0.001 |
| RQ | 1 | 0.00002 | 0.990 | 0.407 |
| EE | 1 | 0.00003 | 1.474 | 0.179 |
| WLR | 1 | 0.00004 | 1.986 | 0.096 |
| body temperature | 1 | 0.00002 | 1.212 | 0.261 |
| sex | 1 | 0.00003 | 1.730 | 0.120 |
| tissue | 4 | 0.02141 | 285.317 | 0.001 |
| model | 11 | 0.02176 | 105.450 | 0.001 |
| residual | 147 | 0.00276 |  |  |

### Consensus gene set

We identified 41 genes that were significantly DE and assigned to at least one significant module for all five tissues as well as were located at least two standard deviations away from the origin in our CCA triplot (Figure 3).

## Discussion

Extensive research has been conducted on the genomic and physiological mechanisms that control water balance in mice (Blumstein and MacManes, 2023; McCue et al., 2017; Rocha et al., 2021). This is particularly intriguing in the context of exploring adaptations to extreme conditions, given that the management of water has significant implications for survival. Studies have largely focused on characterizing the response in kidneys (Rocha et al., 2023; MacManes, 2017; Peng et al., 2023; Rocha et al., 2021), which, while important, represent only a fraction of the physiological and genomic response to dehydration. Indeed, the organismal response to an environmental stressor (such as dehydration), is likely to involve coordination of multiple organ systems and physiological process. However, there are key challenges to studying such responses at the organismal level. Although we understand that organ-systems operate in tandem with other organ systems, the identification of coordination at the level of gene expression is challenging, even when organismal physiology is well-characterized. In addition to complications related to biology, technical complications exist. Here, when looking at gene expression levels across tissues, results highlight differences between the tissues rather than to elucidate the ways in which processes in one depend on the actions of the other, thereby potentially obscuring the gene expression signal of coordination. Further, gene interaction maps (*e.g.,* KEGG) that span organ systems do not currently exist, mostly because the models used for their development (*e.g.,* yeast, flies) lack such complexity present in mammalian systems.

The study of coordination at the level of physiology has been similarly difficult to elucidate, due to limitations in our ability to collect and analyze phenotypic data at a temporal scale that is relevant to the biological phenomenon under study (but see Blumstein et al., 2022; Blumstein and MacManes, 2023; Colella et al., 2021; McKechnie et al., 2021; Ramirez et al., 2022). Despite the challenges, understanding the complex interplay of multi-tissue networked gene expression with whole-organism physiological response is critical to developing a more in-depth understanding of how organisms respond to environmental stressors. Our focus on physiological measurements in a simulated desert environment, as well as on collection of multi-tissue gene expression data in the context of a whole-organism response, represents a distinctive contribution to the field. The findings underscore the complexity of the genetic landscape governing physiological responses to water deprivation and emphasize the need for a broader organismal understanding of the physiological and genomic mechanisms orchestrating successful adaptations.

### Global response to acute water deprivation

The inclusion of physiology and multi-tissue gene expression data in this study provides opportunities to understand how the tissues work together to compensate for water deprivation. During water deprivation, blood volume decreases, which if inadequately managed, may result in a decrease in organ perfusion and ultimately organ failure. The multisystem response to water deprivation involves a coordinated response, including renin-angiotensin-aldosterone system (RAAS) activation, reduced food intake, with compensatory activation of systems for preserving levels of serum glucose, a widespread reduction of genes responsible for clotting factors, and distinct indications of vascular restructuring to support perfusion.

The first response to water deprivation involves the upregulation of processes that are aimed at solute management. In part regulated by the RAAS, this process begins with decreased blood volume detected by baroreceptors found in the carotid sinuses and aortic arch. As a result, the kidneys release the enzyme renin (Blair et al., 1997; Greenleaf, 1992). Renin then acts on a plasma protein called angiotensinogen, produced by the liver, and converts it into angiotensin I (Fountain et al., 2023; Santos et al., 2019). Angiotensin I is subsequently converted into angiotensin II by the angiotensin-converting enzyme (Fountain et al., 2023; Greenleaf, 1992; Santos et al., 2019). Angiotensin II serves as a potent vasoconstrictor, which helps increase tissue perfusion pressure and stimulate the release of aldosterone from the adrenal glands (Fountain et al., 2023; Santos et al., 2019). Aldosterone in turn prompts the kidneys to retain sodium while excreting potassium, which encourages renal water retention. In further support of the activation of the RAAS mechanism, multiple genes in the Renin-Angiotensin system KEGG pathway (hsa04614: Kanehisa et al., 2023; Kanehisa and Goto, 2000) were found to be differentially expressed. Among these genes were *AGT* (angiotensinogen), *ACE2* (angiotensin-converting enzyme), *REN1* (Renin), *CMA1* (converts angiotensin I to angiotensin II), and *AGTR1* (Angiotensin II Receptor) (supplemental table 4), which were all more highly expressed in water-deprived mice. Further, we found two compete pathways of genes to be significantly differentially expressed genes within hsa04614, enhanced vasoconstriction and coagulation cascade as well as vasoconstriction, inflammation, fibrosis, antinatriuresis, reactive oxygen species activation, and Na and water retention (Kanehisa et al., 2023). Interestingly, the candidate gene and essential component of the RAAS mechanism, *AGT,* was significantly upregulated in dehydrated mice, assigned to a significant WGCNA module, and defined as an outlier in the CCA analysis (Table 5). Together with the physiological data, the consistent genetic signature of RAAS activation, using orthogonal analytical methods, suggests a robust whole-body RAAS response which is critical for the cactus mouse to regulate water balance.

**Table 5.** Genes identified in all three analyses (i.e. significantly differentially expressed genes, assigned to a significant module in WGCNA, and are outliers in CCA) in all five tissues in the study (lung (lu), liver (liv), gastrointestinal tract (gi), hypothalamus (hyp), and kidney (kid) grouped by the general function of the gene.

| general function | genes |
| --- | --- |
| housekeeping | ANKRD13D, GPRASP2, ACOX2, ALDOB, GSTA3, HAL, HOGA1, INMT, MTARC1, SPTBN2, DES, EPHX2, FMO5 |
| ion homeostasis | ATP1A2, ALB, APOA4, CYP2E1, CYP4B1, RGN, SLC38A4, AGT, SLC6A13, TF, HP |
| central nervous system | MBP, MAG, PLP1, MAOB, SLC6A13 |
| apoptosis | RELL2, PYCARD |
| coagulation | SERPINF2 |
| blood pressure | ALB, AGT, CASZ1 |
| angiogenesis | ANG, CASZ1 |
| lipid | APOA2, APOA4, APOB, AZGP1, CYP2E1, CYP4B1, SLC27A2 |
| immune response | CFI (renal failure), CTSC, PYCARD, HP |
| gluconeogenesis | CYP2E1 (induced by starvation), PCK1, AGT, SERPINA6 |
| bitter taste | AZGP1 |

At the same time, independent of the RAAS pathway but stimulated by its products, vasopressin is released for the primary function of conserving water. Vasopressin binds to receptors, activating a signaling cascade in the kidneys which functions to retain water by inducing expression of water transport proteins in the late distal tubule and collecting duct, increasing the permeability of the membrane to water. The increased water permeability allows water to move through, from the collecting ducts back into the bloodstream, further aiding in regulating blood volume and fluid balance (Fountain et al., 2023; Santos et al., 2019). Support for the genetic activation of the vasopressin pathway is shown with various genes within the vasopressin-regulated water reabsorption pathway (hsa04962, Kanehisa et al., 2023; Kanehisa and Goto, 2000) are significantly differentially expressed. Specifically, *STX4, RAB5C, RAB11B*, *CREB3L2*, *AQP3*, *VAMP2*, *DYNLL2*, *GNAS*, and *AVPI1* (supplemental table 4) were upregulated in dehydrated mice. Interestingly, *AQP2*, a key gene in hsa04962, did not have any reads mapped to it. It is worth noting here that the identification of differentially expressed transcripts on a KEGG pathway can be thought of in terms of a stoichiometric problem, albeit one where neither current tools nor current data allow us to satisfactorily solve. When attempting to use differential expression to support the upregulation of a given pathway, it may be that only one gene, the quantity of which is rate limiting, may be more highly expressed. Without understanding which transcripts are rate limiting, the interpretation of KEGG pathway mapping may suffer from either over-or under-valuation.

During water deprivation, animals are known to reduce the amount of solid food intake (Armstrong et al., 1980; Hamilton and Flaherty, 1973; Salter and Watts, 2003; Schoorlemmer and Evered, 2002; Watts and Boyle, 2010). Known as dehydration associated anorexia, this secondary response reduces the amount of water required for digestion and facilitates water reabsorption from the kidneys and gastrointestinal tract back to systemic circulation (Rowland, 2007; Watts and Boyle, 2010). While the magnitude of the impact in cactus mice is unknown, diets high in fiber, like the diet consumed in this study, can result in high fecal volume, and has been shown to account for as much as 25% of total daily water loss in rats (Radford Jr, 1959; Rowland, 2007). Perhaps even more importantly, but unquantified, the processing of solid food requires the production of significant quantities of digestive enzymes, all of which are water rich. In humans, as much as 10L of fluids is excreted daily (Ma and Verkman, 1999), which represents a significant investment of water resources. While some of these enzymes continue to be produced during dehydration anorexia, their quantity is likely decreased. As a direct result, the reduction of oral food intake may result in significant water savings, which at least in the short term with the presence of sufficient glycogen stores (see below), should be related to a reduction in water use and electrolyte derangement and therefore enhanced survival.

With limited food intake, there is no external source of glucose, nevertheless, we observed that glucose levels were still maintained (Table 1, Further explained in Blumstein and MacManes, 2023) which is critical for surviving dehydration. This can be achieved by enhancing glycogenolysis and gluconeogenesis (Salter and Watts, 2003; Schoorlemmer and Evered, 2002; Watts and Boyle, 2010), responses that are designed to maintain blood glucose levels during fasting or starvation. In mammals, there is a vasopressin receptor in the liver that when bound activates gluconeogenesis (Bankir et al., 2017), with secondary contributions from the kidney (Nordlie et al., 1999). This suggests a successful two-pronged response to vasopressin secretion, such as vasoconstriction and reabsorption of water from kidney as well as a role in feeding behavior and energy balance. In the current study, we identified the candidate genes *PCK1,* a main control point for the regulation of gluconeogenesis (Hatting et al., 2018) as well as *CYP2E1*, a gene induced by starvation with products involved in gluconeogenesis (Harjumäki et al., 2021; Schattenberg and Czaja, 2014), as upregulated in water-deprived animals, Further, these genes were assigned to a significant module in all five tissues, and defined as an outlier in the CCA analysis (Table 5). Further, the insulin resistance KEGG pathway (hsa04931, Kanehisa et al., 2023; Kanehisa and Goto, 2000) involves various mechanisms contributing to altered glucose metabolism and insulin responsiveness. Notably, all genes but two genes in hsa04931 are significantly up regulated in dehydrated animals (supplemental table 4), all of which are upstream of *PCK1* are in the pathway. Two of these important pathway genes (AGT and SLC27A2, Table 5) were found in our consensus gene set. This upregulation is likely a compensatory response to ensure the maintenance of blood glucose levels in the absence of dietary intake. Lastly, *INMT* (Table 5), a gene in the Tryptophan metabolism pathway (hsa00390, Kanehisa et al., 2023; Kanehisa and Goto, 2000) is upstream of the Glycolysis/gluconeogenesis pathway (map00010, Kanehisa et al., 2023; Kanehisa and Goto, 2000), suggesting complex precursor genes and gene pathways may be contributing to glucose homeostasis as well. The significant changes in expression observed in these genes underscore their role in modulating glucose metabolism, shedding light on how organisms adjust to sustain vital glucose levels during periods of reduced food intake and water deprivation.

During water deprivation, a key challenge is to cope with altered fluid balance and maintain effective blood circulation and nutrient delivery under water-deficient conditions. This process can involve the formation of new blood vessels or the remodeling of existing ones to optimize perfusion, addressing the challenges posed by reduced fluid availability and potential hemoconcentration. We found several overlapping GO terms related to vascular development across all five tissues (vascular development [GO:0001944], circulatory system development [GO:0072359], angiogenesis [GO:0001525], blood vessel development [GO:0001568], blood vessel morphogenesis [GO:0048514], and regulation of vasculature development [GO:1901342]). These GO terms were identified in a myriad of gene modules (water treatment, EE, Hb, TCO2, WLR, and body temperature, Figure 2). Additionally, we identified two genes from our consensus gene set that are mediators of new blood vessel formation and morphogenesis (*ANG*, *CASZ*, Table 5) Prior research has indicated that persistent activation of neuronal systems could modify local blood circulation through angiogenesis. Specifically, rats reared in complex environments had increased capillary density in the visual cortex (Cavaglia et al., 2001). Long-term motor activity has been documented to induce the development of new blood vessels within the cerebellar cortex (Black et al., 1990) and primary motor cortex (Swain et al., 2003), and hyperosmotic stimuli, similar to what experimental animals experience, has been shown to modify the vasculature and induced reversible angiogenesis throughout the hypothalamic nuclei (Alonso et al., 2005). Furthermore, Alim et al. (2019) observed seasonal differences in vascularization, showing blood capillaries were thicker during the winter, suggesting less diffusion across the membrane, compared summer in the dromedary camel.

To maintain blood flow and prevent the formation of clots that could impede circulation, downregulation of coagulation factors during periods of reduced fluid intake could be a protective mechanism against the potential risks associated with increased blood viscosity due to hemoconcentration. As dehydration leads to reduced water content in the blood, resulting in thicker blood consistency there is a risk for blood clot formation. We found several downregulated GO terms related to hemoconcentration (regulation of coagulation [GO:0050818], blood microparticle [GO:0072562], hemopoiesis [GO:0030097], serine hydrolase activity [GO:0017171], and regulation of hemostasis [GO:1900046]) were identified in the hypothalamus and liver (Figure 3) as well as four genes in our consensus gene list involved in blood composition and concentration (*ALB*, *HP*, *SERPINA6*, *SERPINF2*, Table 5). This could indicate several things: 1) To mitigate the impact of hemoconcentration, genes responsible for clotting factors are downregulated to reduce the risk associated with increased blood viscosity as thicker blood is more prone to clotting. 2) to decrease clotting during body temperature dysregulation. Blumstein and MacManes (2023) discuss heterothermy as a mechanism for substantial energy and water savings. In addition to this, relative hypothermia can result in inactivation of coagulation enzymes and/or platelet adhesion defect (Paal et al., 2016), suggesting further mechanisms for maintaining perfusion and limit clotting. However, it’s important to note that the timing of expression poses a potential concern regarding correlations. Decreased body temperature was measured in Blumstein and MacManes (2023) during the dark phase of the experiment and tissues were collected during the light phase of the experiment. Using a single time point snapshot for each experimental condition might overlook certain interactions, given the lag between the expression of a gene (such as a transcription factor) and the expression of downstream effectors.

### Tissue Specific Responses

When analyzing the RNAseq data across all the tissues, we observed that, as expected, samples of the same tissue type clustered together regardless of the water-access treatment, suggesting that the signature of tissues-specific gene expression overpowers the signature of experimental treatment. During individual tissue analysis, the degree of sample separation in PCA space (Supplemental Figure 1) and number of differentially expressed genes suggests tissue specific responses. This is in part supported by downstream gene ontology (GO) terms. Specifically, GO terms related to metabolic processes were identified in multiple significant WGCNA modules in the kidney, but not other tissues. Hydrolase activity and renal system development uniquely in the GI. Lipid metabolic process and detoxification uniquely in the liver. Immune response uniquely in the lung. Lastly, starvation response uniquely in the hypothalamus (Supplemental Figure 2). It is well known that the hypothalamus is the central regulating unit in the brain for maintenance of energy homeostasis (Tran et al., 2022). However, few genes were identified as differentially expressed in the hypothalamus (Figure 1), while our other analyses, WGCNA and CCA, uncovered genes and GO terms that responded to water deprivation. This suggests that the hypothalamus may not be well suited for bulk RNAseq studies due to the heterogeneity of the tissue. Future studies, particularly those using single cell methods (Kephart, 2023; Marquez-Galera et al., 2022; Yue et al., 2023) may further clarify the role that the hypothalamus plays in the overall response to dehydration.

## Conclusion

Here, we highlight the intricate mechanisms involved in regulating water balance in the desert cactus mouse, *P. eremicus*. Our emphasis on whole-organism physiological and multi-tissue (kidney, GI, hypothalamus, liver, and lung) gene expression analysis within a simulated desert environment allowed us to achieve an understanding of genomic mechanisms of water homeostasis. Previous genome scan studies (Colella et al., 2021; Kim et al., 2016; Rocha et al., 2021; Tigano et al., 2020; Wu et al., 2014) have identified many of the same processes we found (i.e., metabolic processes, renal system development, immune response, and starvation response) however, our study design allows us to further interpret function because we were able to both described the tissue specific location and link the processes to physiological measurements.

At a whole-organismal scale, we observed a robust response of the renin-angiotensin-aldosterone system (RAAS) in dehydrated cactus mice, with upregulation of *AGT* in all five tissues as well as upregulation of other pathway genes. Additionally, the compensatory action of *PCK1* was activated in all tissues, further supporting reduced food intake during water deprivation and underscores the body’s adaptive response. However, despite efforts to maintain blood volume, hemoconcentration still occurs, but in response there was a downregulation of genes responsible for coagulation (*e.g., SERPINF2*) as a protective measure against blood clotting in all five tissues, a gene with the major role of regulating the blood clotting pathway. The consequential thickened blood consistency poses challenges to effective blood flow through vessels, compelling the body to initiate the construction of additional vessels to enhance blood movement, further supported by the upregulation of *ANG* in all five tissues, further illustrating the complex interplay of regulatory processes in response to fluid balance disturbances.

## Acknowledgments

We would like to thank members of the MacManes lab for helpful comments and support on previous versions of the manuscript; Adam Stuckert at the University of Huston for lively discussion, valuable insight, and code development; The Animal Resources Office and veterinary care staff at the University of New Hampshire for colony maintenance and care. This work was supported by the National Institute of Health National Institute of General Medical Sciences (R35 GM128843 to M.D.M.).

## Author Contributions

Conceptualization: M.D.M.; Methodology: D.M.B.; Formal analysis: D.M.B., Investigation: D.M.B., Resources: M.D.M.; Writing - original draft: D.M.B.; Writing - review & editing: D.M.B., M.D.M.; Visualization: D.M.B; Supervision: M.D.M.; Project administration: M.D.M.; Funding acquisition: M.D.M.

## Data Availability

https://github.com/DaniBlumstein/dehy_rnaseq

BioProject ID: PRJNA1048512

http://www.ncbi.nlm.nih.gov/bioproject/1048512

---

**Supplemental Figure 1.**
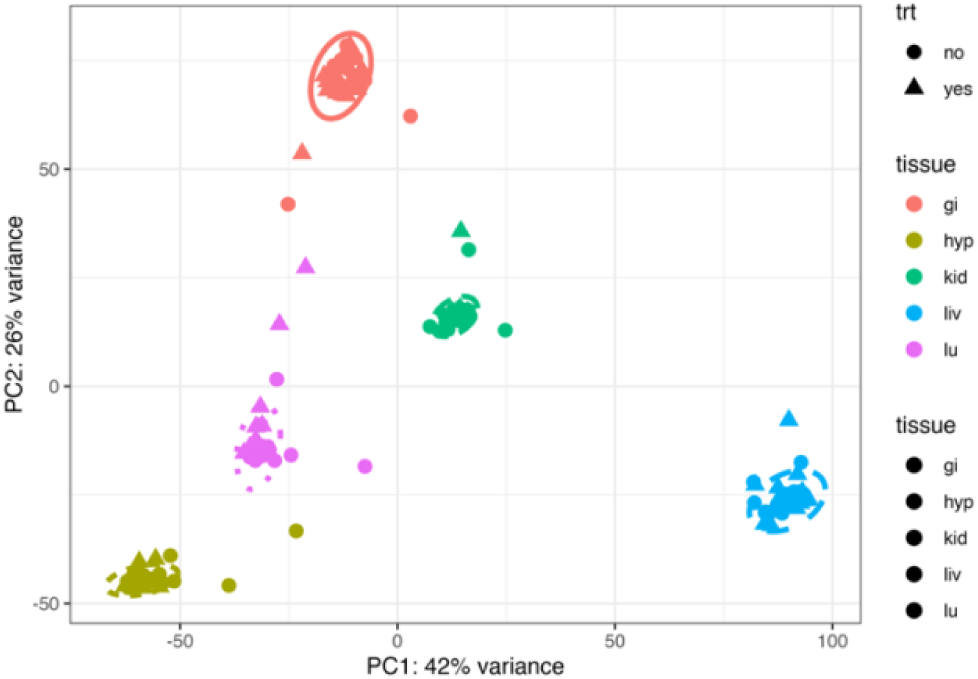
Principal component analysis of gene expression of the lung (lu), liver (liv), gastrointestinal tract (gi), hypothalamus (hyp), and kidney (kid) of *Peromyscus eremicus*. The axes are labelled with the proportion of the data explained by principal components 1 and 2.

**Supplemental figure 2.**
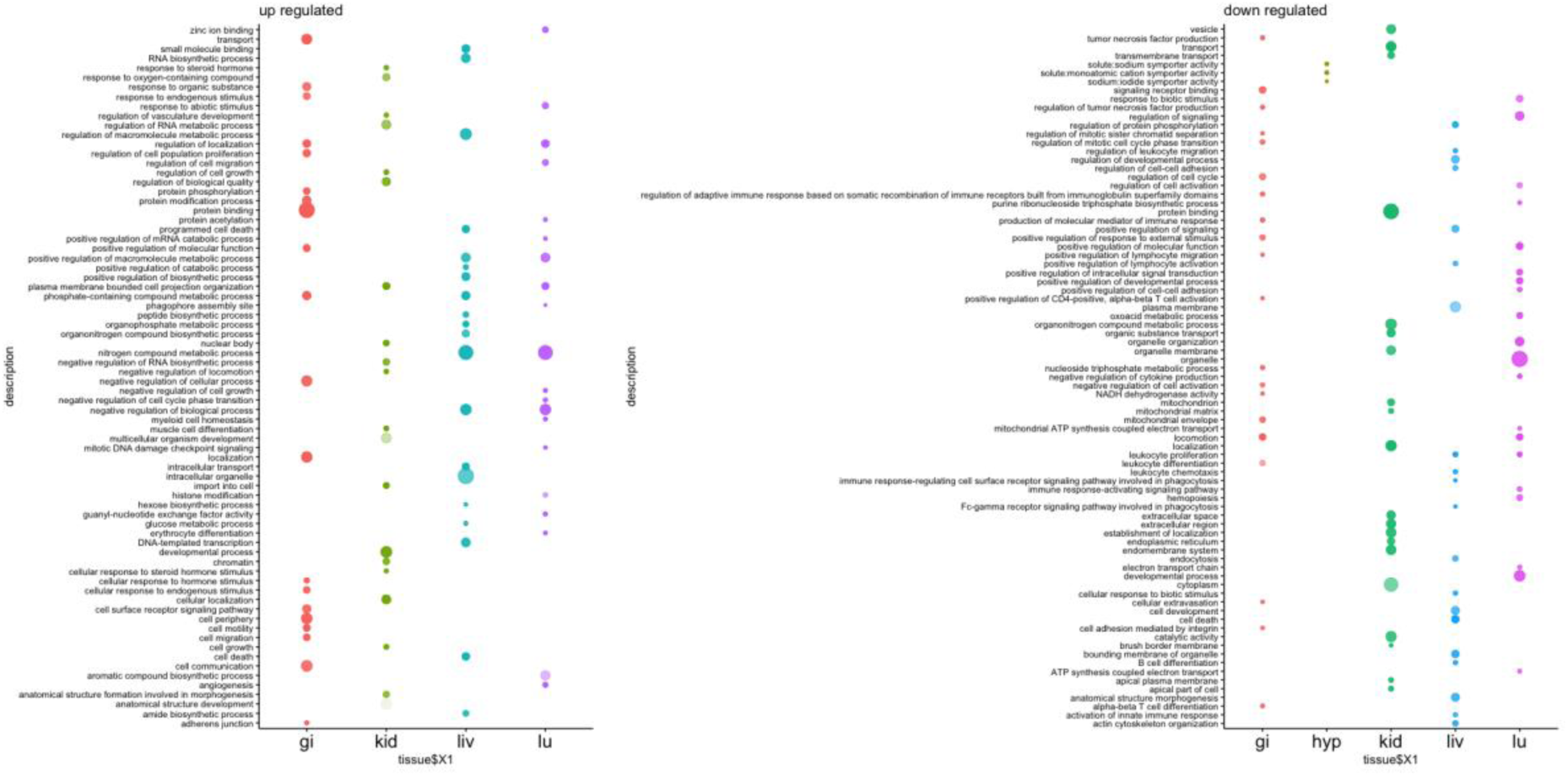
Visualization of gene ontology (GO) terms of up and down regulated differential gene expression between the lung (lu), liver (liv), gastrointestinal tract (gi), hypothalamus (hyp), and kidney (kid) of *Peromyscus eremicus*. Visualized are selections of the top 20 significant GO terms for each phenotype module combination. The number of genes in the GO term are indicated by size of the dots.

**Supplemental table 1.** The number of reads and mapping rate for each RNAseq sample.

**Supplemental table 2.** Gene assignments to WGCNA modules for the lung, liver, gastrointestinal tract, hypothalamus, and kidney of *Peromyscus eremicus*.

**Supplemental table 3.** The number of WGCNA modules for each phenotypic measurement for the lung, liver, gastrointestinal tract, hypothalamus, and kidney of *Peromyscus eremicus*.

| trait | kidney | gastrointestinal tract | liver | lung | hypothalamus |
| --- | --- | --- | --- | --- | --- |
| sex | 0 | 0 | 0 | 1 | 0 |
| delta weight | 5 | 13 | 9 | 7 | 2 |
| proportional weight loss | 7 | 13 | 9 | 8 | 2 |
| Na | 6 | 14 | 12 | 9 | 2 |
| BUN | 7 | 14 | 8 | 8 | 3 |
| AnGap | 1 | 0 | 0 | 0 | 0 |
| K | 1 | 0 | 0 | 0 | 0 |
| Cr | 2 | 3 | 2 | 5 | 4 |
| Htc | 7 | 12 | 10 | 8 | 3 |
| Cl | 6 | 12 | 12 | 7 | 2 |
| Glu | 0 | 0 | 0 | 0 | 0 |
| Hb | 7 | 12 | 10 | 8 | 3 |
| TCO2 | 5 | 9 | 10 | 6 | 2 |
| iCa | 2 | 0 | 4 | 2 | 0 |
| RQ | 5 | 1 | 1 | 1 | 2 |
| EE | 0 | 0 | 2 | 1 | 1 |
| WLR | 4 | 3 | 6 | 6 | 1 |
| body temperature | 2 | 3 | 9 | 5 | 2 |
| water access | 7 | 12 | 9 | 7 | 1 |
| Total modules | 12 | 24 | 18 | 13 | 18 |

**Supplemental table 4.** Results, including the log fold change values, p values, and adjusted p values, from the differential gene expression analysis for for the lung (lu), liver (liv), gastrointestinal tract (gi), hypothalamus (hyp), and kidney (kid) of *Peromyscus eremicus*.

