## Supplementary figures and images for "When the tap runs dry: The multi-tissue gene expression and physiological responses of water deprived *Peromyscus eremicus*"

### Supplemental figure 1

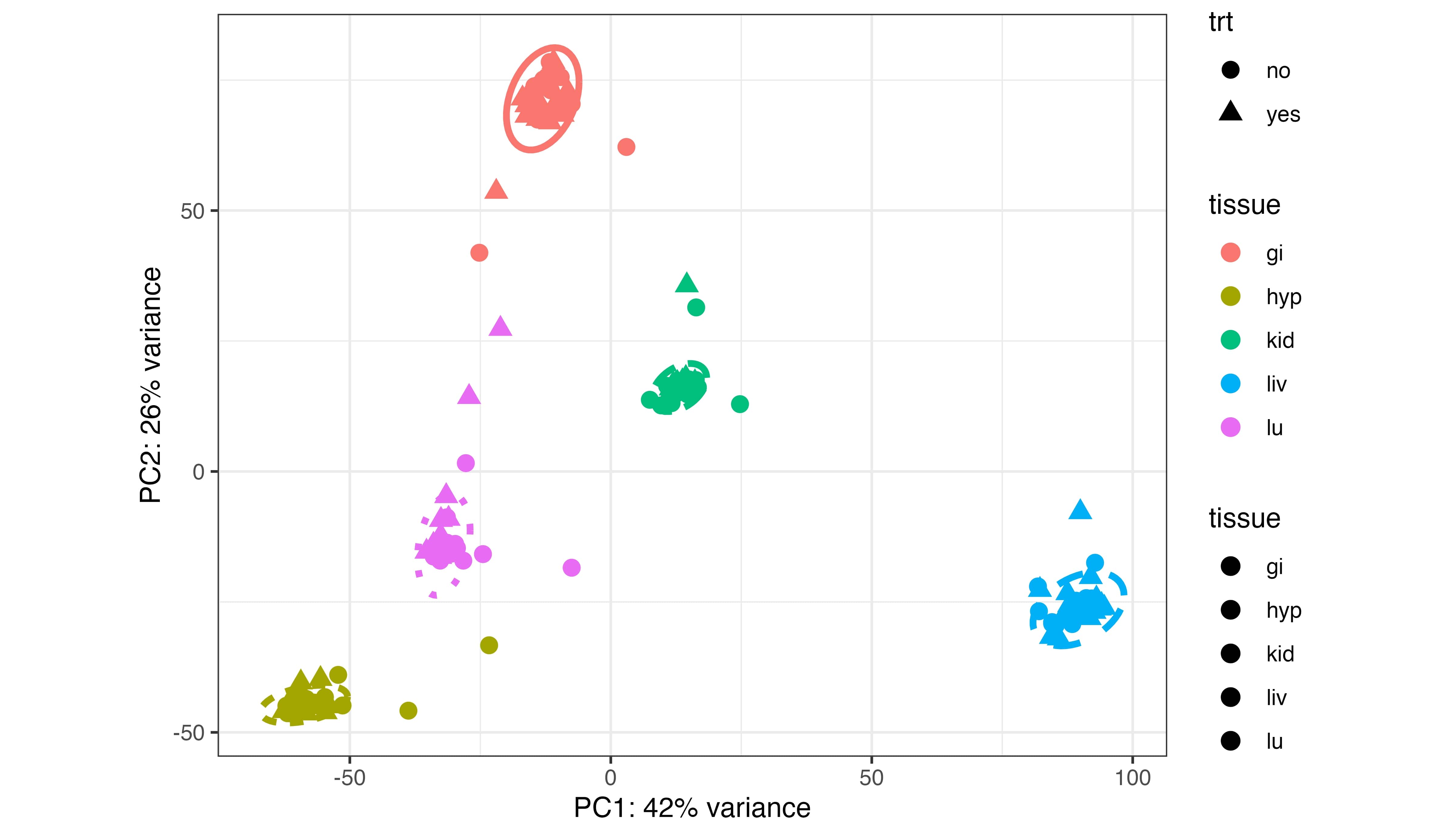

### Supplemental figure 2

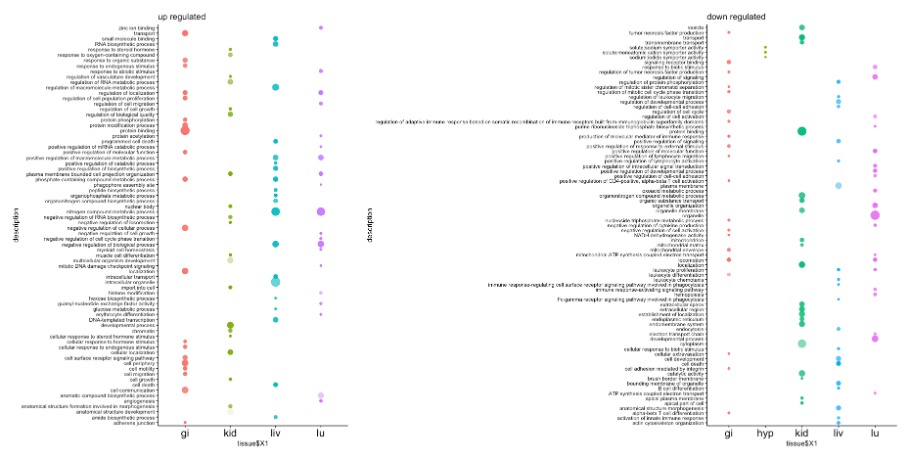
